# Spatially Resolved Microglial Expression Around Aβ Plaques in Human Alzheimer’s Disease Tissue

**DOI:** 10.1101/2025.08.29.673089

**Authors:** Jack I. Wood, Georgia Ppasia, Paul Rolland-Du-Roscoat, Sneha Desai, Shamim Choudhury, Aya Balbaa, Daria Gavriouchkina, Modesta Blunskyte-Hendley, John Hardy, Damian M. Cummings, Dervis Salih, Jörg Hanrieder, Frances A. Edwards

## Abstract

Using microglia-enriched spatial transcriptomics on human Alzheimer’s disease tissue, we identify distinct gene expression changes across microglia located in direct contact with plaques, in periplaque regions, and in areas distant from plaques. We define a group of plaque contact–only microglial (PCOM) genes whose expression increases exclusively in microglia directly contacting plaques. These genes show significant overlap with previously reported gene sets, suggesting that many of the well-characterised disease-associated microglia (DAM) and other AD-related gene-expression signatures are only upregulated when microglia contact plaques. We further identify distinct co-expression networks associated with disease-relevant covariates, including an immune module linked to APOE genotype and a synaptic–mitochondrial module negatively associated with Braak stage. Finally, we compare the human dataset to our previously published data from 18-month-old *App^NL-F^* mice, generated using the same experimental paradigm and demonstrate cross-species concordance in gene expression particularly within plaque-contacting microglia.

## Introduction

The immune response to Alzheimer’s disease (AD) pathology has been consistently implicated as a major contributor to both AD risk and disease progression. Genome-wide association studies (GWAS) have identified more than 75 genetic loci associated with AD, many of which show preferential expression in microglia, the brain’s main innate immune cell (Bellenguez et al., 2022; Wightman et al., 2021). These genetic findings strongly support an essential role for microglia in AD, aligning with phenotypic observations of their clustering around Aβ plaques (Condello et al., 2015; Itagaki et al., 1989).

The advent of microarray-based transcriptomic profiling enabled unbiased, genome-wide analysis using mRNA-specific probes. Early microarray studies of human brain tissue revealed the upregulation of immune signalling pathways, including cytokines and chemokines, pointing to a significant microglial inflammatory component in AD pathogenesis (Weeraratna et al., 2007; Xu et al., 2006). Later, analyses in AD mouse models confirmed widespread microglial transcriptional changes, identifying immune-associated gene modules that correlated strongly with Aβ plaque pathology (Matarin et al., 2015). The second major advance came with bulk RNA sequencing, which overcame reliance on probe sets and expanded the resolution of microglial networks. Bulk sequencing studies reinforced the link between plaques and innate immune activation, revealing co-expressed gene sets involving complement components, lysosomal machinery, and multiple AD risk loci (Humphries et al., 2015; Salih et al., 2019). Together, these findings established that microglial responses to amyloid pathology are highly coordinated and involve diverse cellular processes extending beyond classic inflammatory signalling. Single-cell RNA sequencing then provided a third breakthrough by enabling the isolation and profiling of individual microglia. This approach revealed striking heterogeneity within the microglial population, distinguishing homeostatic cells from subtypes that emerge in the context of AD pathology (Keren-Shaul et al., 2017; Sala Frigerio et al., 2019). Among these, disease-associated microglia (DAM), also known as activated response microglia (ARM), were defined by a transcriptional programme enriched for genes involved in phagocytosis, lipid metabolism, and immune signalling. Subsequent studies confirmed the existence of multiple microglial states across mouse and human AD tissue, highlighting the dynamic and context-dependent nature of microglial responses (Grubman et al., 2021; Olah et al., 2020; Sun et al., 2023).

Although it had been shown that these AD associated states were closely correlated with plaque density (Matarin et al., 2015), their spatial relationship to pathology could not be established without direct tissue-resolved methods. The fourth advance, spatial transcriptomics, enabled gene expression to be profiled within defined tissue regions. This led to the identification of plaque-induced genes (PIGs), whose expression increased in proportion to local plaque density within the same animal (Chen et al., 2020). These findings provided the first transcriptome-wide evidence linking Aβ pathology directly to the enrichment of immune and lysosomal pathways in surrounding tissue.

We previously extended the analysis of plaque-induced genes (PIGs) by resolving their spatial enrichment in relation to Aβ plaques in aged *App^NL-F^* mice using spatial transcriptomics. This revealed that many AD-associated genes, including complement components, cathepsins, and *Trem2*, showed increased expression only in plaque-contacting microglia (Wood et al., 2022). The development of improved mouse models of AD, such as the *App^NL-F^* model, has been driven by the historical lack of successful translation from mouse studies to human clinical trials (Drummond and Wisniewski, 2017), highlighting the need for continuous improvement of animal models and validation with associated findings in human tissue.

In the present study, we use spatial transcriptomics to analyse microglial gene expression in select regions of human AD hippocampal tissue: microglia directly contacting plaques, those in periplaque regions, and those distant from plaques, the same experimental paradigm as Wood et al. (2022). We demonstrate that genes upregulated exclusively in plaque-contacting microglia show significant enrichment for disease-associated microglia (DAM) profiles and other AD-related gene sets.

Furthermore, we identify distinct gene modules associated with Braak stage and APOE genotype. Finally, by directly comparing human AD tissue with our previous *App^NL-F^* mouse dataset, we reveal a conserved transcriptional response in plaque-contacting microglia across species.

## Methods

### Human Tissue: Licences and Ethics

The brain samples and samples were obtained from The Netherlands Brain Bank, Netherlands Institute for Neuroscience, Amsterdam (open access: www.brainbank.nl). All Material has been collected from donors for or from whom a written informed consent for a brain autopsy and the use of the material and clinical information for research purposes had been obtained by the NBB.

The project was approved by the UCL ethics committee, Ethics ID: 26399.001. Transfer from the Netherlands Brain Bank under the Material Transfer Act, Project 1612. The storage of FFPE human brain tissue was with the UCL dementia research institute, HTA licence: 12198. The case summary of all tissues used for this project is found in Table 1.

**Table 1.** Summary of human hippocampal sample characteristics.

| Clinical Diagnosis | Plaque Positivity | Group | *APOE* Status | Age at Death | Sex | Year of Death | Post Mortem Delay (hrs:min) | Brain Weight (g) | Braak Stage |
| --- | --- | --- | --- | --- | --- | --- | --- | --- | --- |
| AD | + | AD | 43 | 94 | m | 2015 | 04:15 | 945 | 4 |
| AD | + | AD | 33 | 78 | f | 2016 | 04:55 | 1130 | 6 |
| AD | + | AD | 33 | 94 | m | 2016 | 04:15 | 870 | 5 |
| AD | + | AD | 33 | 96 | f | 2016 | 04:15 | 940 | 5 |
| AD | + | AD | 33 | 87 | m | 2016 | 05:30 | 1045 | 5 |
| AD | + | AD | 33 | 73 | f | 2017 | 04:10 | 970 | 6 |
| AD | + | AD | 33 | 96 | m | 2017 | 04:25 | 1095 | 5 |
| AD | + | AD | 43 | 81 | f | 2017 | 05:30 | 1015 | 6 |
| AD | + | AD | 43 | 87 | f | 2017 | 03:25 | 1193 | 4 |
| AD | + | AD | 33 | 86 | m | 2017 | 04:50 | 1295 | 4 |
| AD | + | AD | 43 | 76 | m | 2018 | 04:40 | 1300 | 6 |
| AD | + | AD | 43 | 86 | m | 2018 | 03:45 | 1406 | 4 |
| AD | + | AD | 43 | 68 | m | 2018 | 04:30 | 1163 | 6 |
| AD | + | AD | 43 | 97 | m | 2018 | 05:10 | 1120 | 5 |
| AD | + | AD | 33 | 95 | m | 2020 | 06:05 | 1245 | 5 |
| AD | + | AD | 43 | 106 | f | 2020 | 04:50 | 1100 | 4 |
| AD | + | AD | 43 | 83 | f | 2021 | 04:30 | 1155 | 4 |
| AD | + | AD | 33 | 88 | f | 2021 | 05:55 | 1040 | 5 |
| AD | + | AD | 43 | 73 | f | 2021 | 05:15 | 1055 | 6 |
| ND | + | CU-AP | 43 | 82 | m | 2017 | 05:45 | 1195 | 2 |
| ND | + | CU-AP | 33 | 101 | f | 2019 | 06:00 | 896 | 3 |
| ND | + | CU-AP | 43 | 79 | f | 2020 | 06:00 | 1075 | 3 |
| ND | + | CU-AP | 43 | 84 | f | 2016 | 07:40 | 1165 | 3 |
| ND | + | CU-AP | 33 | 83 | m | 2016 | 05:05 | 1108 | 2 |
| ND | + | CU-AP | 43 | 103 | f | 2018 | 09:50 | 1155 | 3 |
| ND | + | CU-AP | 43 | 73 | m | 2014 | 04:25 | 1285 | 1 |

### Human Tissue Samples

FFPE hippocampal tissue samples were obtained from posterior, mid, and anterior regions of the hippocampus, according to tissue availability. Tissue samples presenting with mixed pathologies were not chosen to enable clearer interpretation of Aβ plaque–specific changes and to minimise potential off-target binding of Amytracker to other β-pleated sheet–rich pathologies. Whole brain tissue was fixed in formalin for 4 weeks before individual regions were dissected and embedded in paraffin. Clinical AD diagnosis was initially based on the criteria established by McKhann et al. (1984) and later updated by Dubois et al. (2007) for probable AD. The Netherlands Brain Bank performed the final confirmation of AD by histopathological assessment.

### Tissue Sectioning

FFPE blocks were sectioned at 8 μm using a rotary microtome. Three of these tissue squares were mounted per SuperFrost Plus slide within the designated GeoMx scan area.

### GeoMx Slide preparation

Slide preparation followed the standard *GeoMx Slide Preparation* protocol with minor optimisations. Slides were baked at 60 °C for 1 h, paraffin was removed in xylene, followed by a 100 % ethanol wash. Sections were rehydrated through 95 % ethanol in 0.1 M PBS. For antigen retrieval, slides were incubated in 10 mM Tris and 1 mM EDTA for 35 min at 100 °C in a slide steamer. RNA retrieval was performed using minimal protein digestion with 0.5 µg/ml proteinase K in 0.1 M PBS for 15 min at 37 °C. Sections were fixed in 4 % formaldehyde and quenched with 0.1 M glycine and 0.1 M Tris in DEPC-treated water. Whole transcriptome probe hybridisation was carried out by immersing sections in probe mix (per slide: 225 µl Buffer R, 25 µl GeoMx Whole Transcriptome Probe Mix, 25 µl DEPC-treated water) and incubating overnight at 37 °C in a hybridisation oven (HybEZ^TM^ II, #321720). Non-hybridised, weakly hybridised, and non-specifically bound probes were removed with 50 % formamide in 2× SSC twice for 25 min at 37 °C in a water bath. Non-specific binding was blocked in Buffer W for 30 min at RT. Sections were then incubated with primary antibodies (rabbit anti-IBA1, 1:300; FujiFilm Wako Chemicals, #019-19741) diluted in Buffer W for 4 h at RT, followed by secondary antibodies (goat anti-rabbit 594, 1:500; Thermo Fisher Scientific, #A11037) in Buffer W for 2 h at RT. Amyloid plaques were stained with Amytracker 520 dye (1:1000 in 0.1 M PBS; Ebba Biotech) for 30 min at RT. Slides were immersed in 2× SSC and stored overnight at 4 °C.

### GeoMx ROI/AOI selection and barcode collection

One section from each human hippocampus was analysed, with 2–4 technical replicates per AOI type per section collected. ROIs and AOIs were defined based on morphological stains.

Regions of interest (ROIs):

1. A region drawn around an area of heavy plaque load in which all microglia would fall within 50 μm of a plaque. This was only performed in plaque-bearing tissue.
2. A region drawn which did not contain plaques and with borders >100 μm from any visible plaque. This was performed on all tissues.

Areas of illumination (AOIs):

a. Within ROI1 - “Plaque Contacting Microglia”, co-localisation of Amytracker520 and IBA1;
b. Within ROI1 - “Periplaque Microglia”, IBA1 signal in ROI 1 not colocalised with Amytracker520 and
c. Within ROI2 - “Away Microglia”, all IBA1 signal in ROI 2.

Since tau tangles also exhibit beta-sheet morphology, Amytracker stained both plaques and tau tangles. However, these pathologies are often regionally separated; therefore, areas of tangle positivity were avoided to enrich for Aβ plaque-associated effects only.

### GeoMx Library preparation

AOI aspirates were dried for 1 h at 60 °C in a thermocycler and subsequently rehydrated in 10 µl RNase-free water. PCR reactions were prepared in each well containing 4 µl of the corresponding well from the GeoMx DSP rehydrated collection plate, 4 µl of the matching well from the Seq Code primer plate, and 2 µl GeoMx master mix. The plates were thermocycled as follows: 30 min at 37°C, 20 min at 50°C, 3 min at 95°C, 18 cycles of [15 s at 95°C, 60 s at 65°C, 60 s at 68°C], 5 min at 68°C, ending at 4°C. PCR products were pooled. To remove impurities, the PCR product was DNA purified with two cycles of AMPure XP bead purification. Each cycle included: addition of AMPure XP beads for 5 min, pellet formation on a magnetic stand, supernatant removal, two washes with 80% ethanol (discarding ethanol each time), and resuspension in elution buffer. A D1000 TapeStation system was employed to confirm the successful collection and preparation of GeoMx DNA oligomer barcode libraries. RNA sequencing of the barcode-derived cDNA libraries was carried using the Illumina NextSeq 2000 with X-LEAP chemistry.

### Probe quality control

FASTQ files were converted to .DCC files using the GeoMx NGS GUI. DCCs containing barcode counts were translated to gene symbols with the human probe assay metadata PKC files. Counts were matched to their associated AOIs with the sample annotation file provided from the GeoMx collection run. AOIs that did not meet a minimum number of 1000 reads per AOI, ≥80% of reads in an AOI successfully trimmed, ≥80% of reads in an AOI successfully stitched, ≥75% of reads in an AOI aligned to the reference, and ≥50% sequencing saturation were removed from further analysis. Raw GeoMx DSP counts were processed in R (v4.3.1). Low-signal samples and AOIs were excluded based on library size per area distributions. Genes with expression below the 75th percentile of negative probes were removed.

### Mouse data

The data presented in this study on *App^NL-F^* mice were extracted and re-analysed from a previous study, Wood et al. (2022).

### Differential expression analysis

Raw count data from both mouse and human experiments were processed using a consistent pipeline. Imported counts were loaded into R and converted into a DGEList object using edgeR. The design matrix included AOI type (plaque, periplaque, away) as the main factor, with APOE status, sex, diagnostic group, and batch incorporated as covariates in the human analysis. Counts were transformed with precision weights using the voom method in limma (Ritchie et al., 2015). To account for multiple AOIs sampled per subject, a consensus correlation was estimated with duplicateCorrelation. Linear models were fitted for each gene, and empirical Bayes moderation was applied. Differentially expressed genes were identified using an adjusted P value < 0.05 (Benjamini–Hochberg FDR).

### Analysis pipelines

All analyses were performed in R version 4.3.1:

#### Gene Ontology

Gene Ontology enrichment analysis was performed using the clusterProfiler package (Yu et al., 2012). Gene-sets were converted to Entrez IDs and tested for over-representation in the Biological Process database using the enrichGO function with the org.Hs.eg.db database. Benjamini–Hochberg method was used for multiple testing correction, with significance thresholds of q < 0.05. Semantic similarity between Gene Ontology (GO) biological process terms was calculated using the GOSemSim R package, which applies ontology-based measures (e.g. Wang’s method) to quantify functional relatedness (Yu, 2020). Similarity scores range from 0 (no relation) to 1 (identical terms).

#### WCGNA

Voom-transformed counts were analysed using the WGCNA package (Langfelder and Horvath, 2008). Modules were detected via dynamic tree cutting on the topological overlap matrix. Module eigengenes were correlated with ROI type, APOE, sex, diagnosis, and Braak stage using Pearson correlations and p-values determined by linear mixed models accounting for repeated measures. Intramodular connectivity, and networks were exported to Cytoscape (v3.10.3) for visualisation. Top 50 nodes shown per module based on connectivity, where available. Modules with fewer than 50 nodes above the edge threshold (0.08) are shown in full.

#### Cell type enrichment

Module genes were compared to reference single-cell RNA-seq profiles from human cortex (Zhang et al., 2016). Replicates within each cell type (Neurons (N), astrocytes fetal (AF), astrocytes mature (AM), microglia (M), oligodendrocytes (O), endothelial (E)) were averaged. Genes with zero variance across cell types were removed, then expression was row-wise z-scored. Heatmaps were generated with pheatmap and clustered by complete linkage; enrichment was defined as the cell type with the highest relative expression.

#### GSEA

Mouse genes were converted to human orthologs using the babelgene package. Gene set enrichment analysis (GSEA) was performed with clusterProfiler using mouse-analysis-derived gene sets as TERM2GENE against human rankings.

#### RRHO

Threshold-free cross-species concordance was evaluated using the RRHO2 package (Cahill et al., 2018). Mouse genes were mapped to human orthologs, and both species were ranked by logFC. Overlap enrichment across all rank thresholds was quantified via hypergeometric testing with Benjamini–Yekutieli correction (stepsize = 10), and concordance was visualised as −log10(p) heatmaps. For visualisation, we truncated extreme significance by capping −log10(P) at 30 to stabilise the heatmap colour scale.

#### Venn diagrams

Venn diagrams were generated with ggVennDiagram and VennDiagram packages. Enrichment of overlapping genes was quantified using one-sided Fisher’s exact tests with a total background of all tested genes.

#### Transcription factor analysis

High-confidence human TF–target interactions (confidence A–C) were obtained from DoRothEA and converted into VIPER-compatible regulons. TF activity scores were inferred from the voom-normalised expression matrix using VIPER (Alvarez et al., 2016; Garcia-Alonso et al., 2019). Differential TF activity between AOI types (Plaque, Periplaque, Away) was tested using limma and accounting for repeated measures via duplicateCorrelation. Multiple testing correction was performed using the Benjamini–Hochberg method.

#### PEA plot

Evidence for neuroprotective or neurotoxic effects were compiled from peer-reviewed studies investigating the selected gene. Each study contributed a score of +1 to either the neuroprotective or neurotoxic category based on the reported impact. Studies without a clearly defined direction of effect were excluded.

### Code availability

The code and associated data directory have been deposited in Zenodo and are accessible via DOI: 10.5281/zenodo.16947474.

### Data Availability

Data can be explored through the interactive online database at https://edwardslab.shinyapps.io/Humaneac/. Raw data have been deposited in NCBI GEO under accession number GSE306381.

## Results

### ROI selection for microglia-enriched spatial transcriptomics on human tissue

The experimental design closely parallels our previous study in 18-month-old *App^NL-F^* mouse models of AD (Wood et al., 2022). Here, the observation that many AD-associated microglial genes only increase in expression upon microglial-plaque contact could be an artefact of using AD mouse models. To assess the translatability of this finding, we performed GeoMx spatial transcriptomics on *postmortem* human hippocampal tissue. An antibody against IBA1 was used to label microglia, and the beta-sheet dye Amytracker was applied to detect Aβ plaques (Figure 1A). Note that this is a more conservative definition of plaques than used in our mouse study, where we defined plaques as all amyloid beta antibody positive pixels hence including the diffuse plaques and the halo around the structured beta-pleated regions that are labelled by Amytracker. Guided by the immunofluorescence images, gene expression was profiled within selected regions of interest (ROIs). Within each ROI, areas of illumination (AOIs) defined the thresholded regions from which transcriptomic data were collected. These were: microglia directly contacting plaques (Amytracker and IBA1+ve pixels), microglia in the periplaque region (IBA1+ve, Amytracker-ve pixes in plaque regions), and microglia located away from plaques (Figures 1B & 1C).

**Figure 1.**
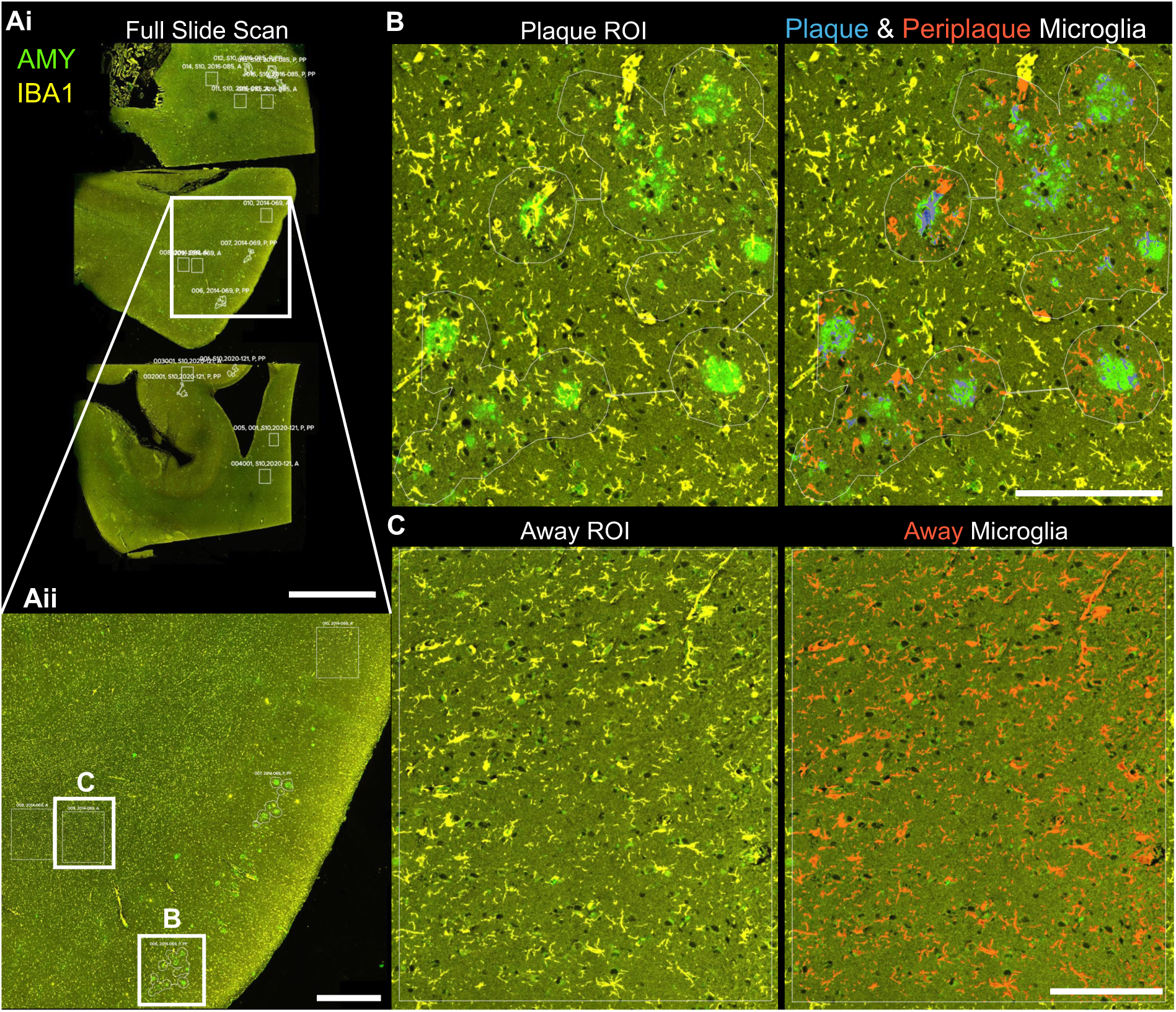
ROI selection for microglia-enriched spatial transcriptomics on human tissue. Representative staining and region of interest (ROI) selection for spatial transcriptomics analysis. 8 μm-thick sections were labelled for microglia (IBA1 antibody, yellow) and Aβ plaques (Amytracker, green).(Ai) Full-slide scan showing three hippocampal sections. Scale bar: 5000 μm. (Aii) Zoomed-in image of the highlighted region in (Ai). Scale bar: 1000 μm. (B) Example of a plaque ROI. The right panel highlights the areas collected for ‘plaque’ (blue) and ‘periplaque’ (red) microglial regions of interest. Scale bar: 200 μm. (C) Example of an ‘away’ microglia ROI. The right panel shows the area collected for ‘away’ microglia (red). Scale bar: 200 μm.

Unlike for mouse models, in human AD tissue, factors affecting inter-sample variation, such as genetics, environment, age at death, and postmortem delay, cannot be controlled. To minimise inter-sample variation while preserving RNA integrity, FFPE hippocampal tissues were selected for a short postmortem delay and death having occurred within the last decade. Samples were selected to achieve a balanced distribution by *APOE* status and sex (Table 1). A key advantage of spatial transcriptomics in overcoming inter-sample variability is the fact that the different types of AOI can be compared within the same tissue section.

### Microglial plaque contact is necessary for AD-associated gene expression in Alzheimer’s disease

GeoMx counts were analysed using the limma-voom pipeline, incorporating APOE status, diagnostic group, and sex as covariates in the model to isolate Aβ plaque– specific changes in gene expression. To visualise the global transcriptional effects of these covariates across the dataset, volcano plots of the main effects of group, sex, and *APOE* status from the limma model are presented in Figure S3.

Based on plaque proximity, three pairwise comparisons were made: plaque-contacting vs away microglia (Figure 2Ai), plaque-contacting vs periplaque microglia (Figure 2Aii), and periplaque vs away microglia (Figure 2Aiii). As all AOIs were extracted from microglia, PCA revealed only small but distinct changes across AOIs (Figure S1). The expression profiles of individual analysed genes can be explored interactively at https://edwardslab.shinyapps.io/Humaneac/. Complete AOI comparisons can be found in Supplemental Table 1.

**Figure 2.**
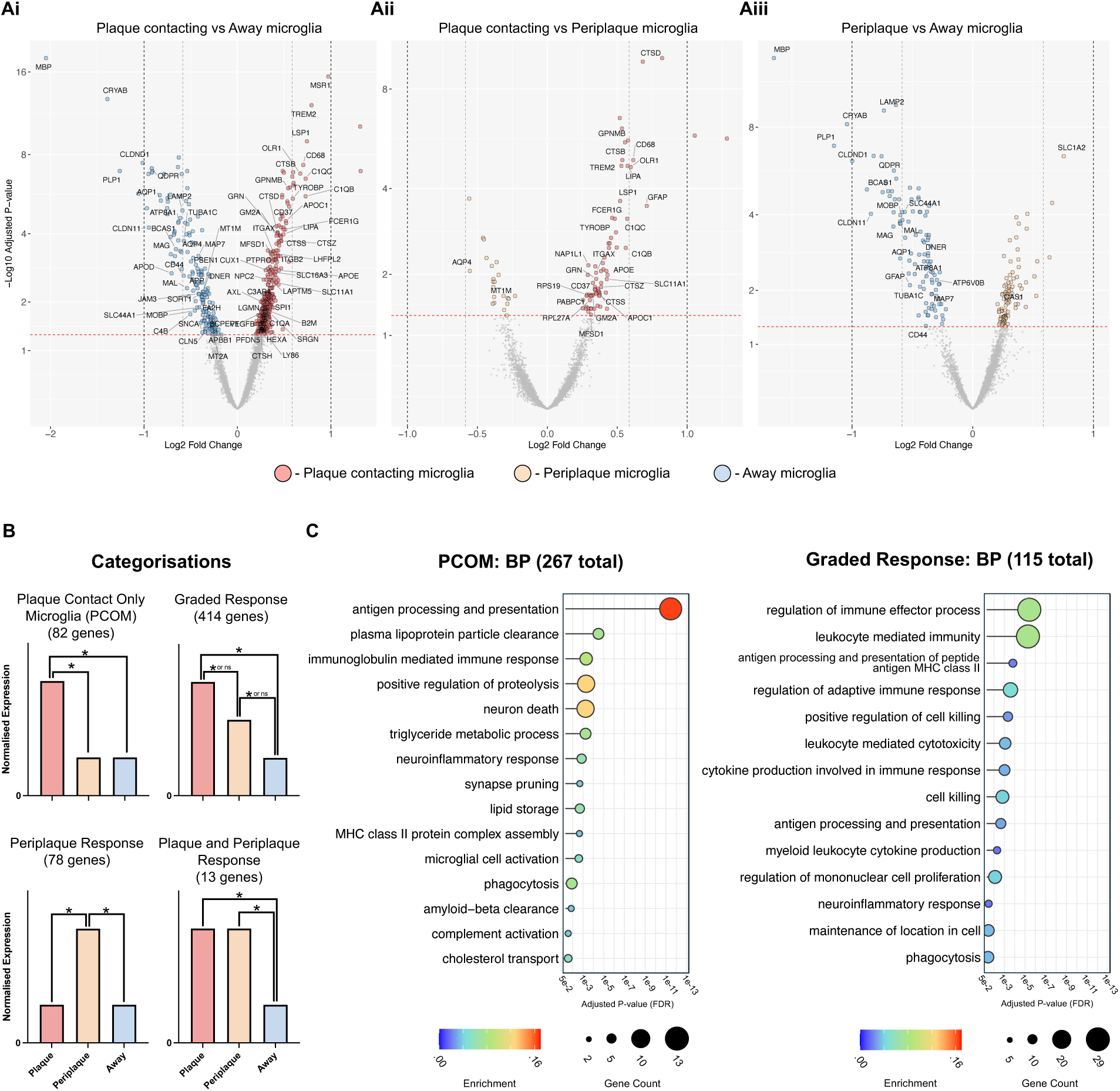
Microglial plaque contact is necessary for AD-associated gene expression in Alzheimer’s disease. (A) Volcano plots showing differential expression analyses for microglia in (Ai) Plaque vs Away, (Aii) Plaque vs Periplaque, and (Aiii) Periplaque vs Away regions. (B) Representative categories illustrating different gene expression responses to plaques, with example graphs for each category. (C) Gene ontology analysis of PCOM genes, highlighting select enriched terms from the Biological Process database. The total number of significant terms identified in each database is indicated in brackets. Statistical analysis: (A) limma-voom with covariate adjustment and adjusted for repeated measures. (C) P values were were identified using the enrichGO() function from clusterProfiler and calculated using a hypergeometric test. (A & C) P values are adjusted for multiple comparisons using Benjamini-Hochberg false discovery rate (FDR) method. Human cases (n = 26).

As in our previous spatial transcriptomics analysis of *App^NL-F^* mice (Wood et al., 2022), the expression patterns across the three AOIs allowed each gene to be classified into one of four categories (Figure 2B):

- Plaque-Contact Only Microglial genes (PCOM): increased expression exclusively in microglia directly contacting plaques;
- Graded Response Microglial genes: a stepwise increase in expression from away to periplaque microglia, and further from periplaque to plaque microglia;
- Periplaque-Responsive Microglial genes: increased expression enriched in microglia within the periplaque region compared to other regions and
- Plaque and Periplaque-Responsive Microglial genes: increased expression in both periplaque and plaque microglia relative to away microglia, with no significant difference between the two.

The equivalent categories were also created for downregulated genes (Figure S2). Supplemental Table 2 contains the gene lists associated with each category.

Gene enrichment analysis was performed for each category using the Biological Process branch of the Gene Ontology database (Figure 2C). The 82 PCOM genes enriched terms associated with an immune response, driven by genes such as *SPP1*, *CTSD*, *C1QC*, and *C1QB* (Figure 2C). Consistent with findings from Wood et al. (2022), we also again observed a plaque-contact-specific increase in *TREM2* expression. Antigen processing and presentation pathways were also overrepresented in PCOM, with contributions from multiple HLA genes including *HLA-B*, *HLA-DOB*, *HLA-DRB1*, and *HLA-DPB1*. Additionally, genes involved in lipid storage, such as *APOE*, *GM2A*, and *FITM2*, were significantly enriched in this category. The graded response category, comprising 414 genes, was similarly enriched for immune-related terms such as regulation of immune effector processes and neuroinflammatory response (Figure 2C). These genes tended to be related to neuroinflammatory processes, including those involved in cytokine signalling pathways, such as *IL13*, the TNF-α receptor *TNFRSF1B*, and inflammasome-associated genes including, *PYCARD*, *P2RX7*, and *MYD88*. The remaining two gene categories did not show significant enrichment for terms within the biological process ontology suggesting that most of the responses of microglia are either dependent on the direct contact with plaques or, if upregulated in the immediate vicinity of plaques, the upregulation becomes increasingly strong as they come closer to plaques.

Genes downregulated specifically in plaque-contacting microglia were enriched for only two terms, both related to the response to metal ions, driven by the expression of metallothionein-encoding genes *MT1M* and *MT2A* (Figure S1). In contrast, genes downregulated across both plaque and periplaque regions were significantly enriched for myelin-related terms, including *MBP*, *PLP1*, and *MAG*. All enriched terms and associated genes can be found in Supplemental Table 3. An important limitation of GeoMx spatial transcriptomics is that, although barcode capture is guided by 2D images, transcripts are extracted from the underlying 3D tissue, resulting in potential contamination from cell types located above or below the targeted microglia. This seems to come up particularly in the genes found to be downregulated in the vicinity of plaques.

### PCOM genes are significantly enriched for the DAM expression profile

The gene expression response of microglia to AD-associated pathology has been consistently reported across multiple studies, including their enrichment among genes located at loci containing AD risk-associated SNPs. To investigate the spatial specificity of such responses, PCOM and graded response genes were tested for enrichment against several well-established AD-associated microglial gene sets (Figure 3A). Here, PCOM genes showed significant enrichment across all tested gene sets, whereas graded response genes failed to significantly enrich the majority of these gene sets, including the widely cited DAM signature. This supports the interpretation that these canonical AD-associated microglial signatures predominantly reflect microglia in direct contact with plaques.

**Figure 3.**
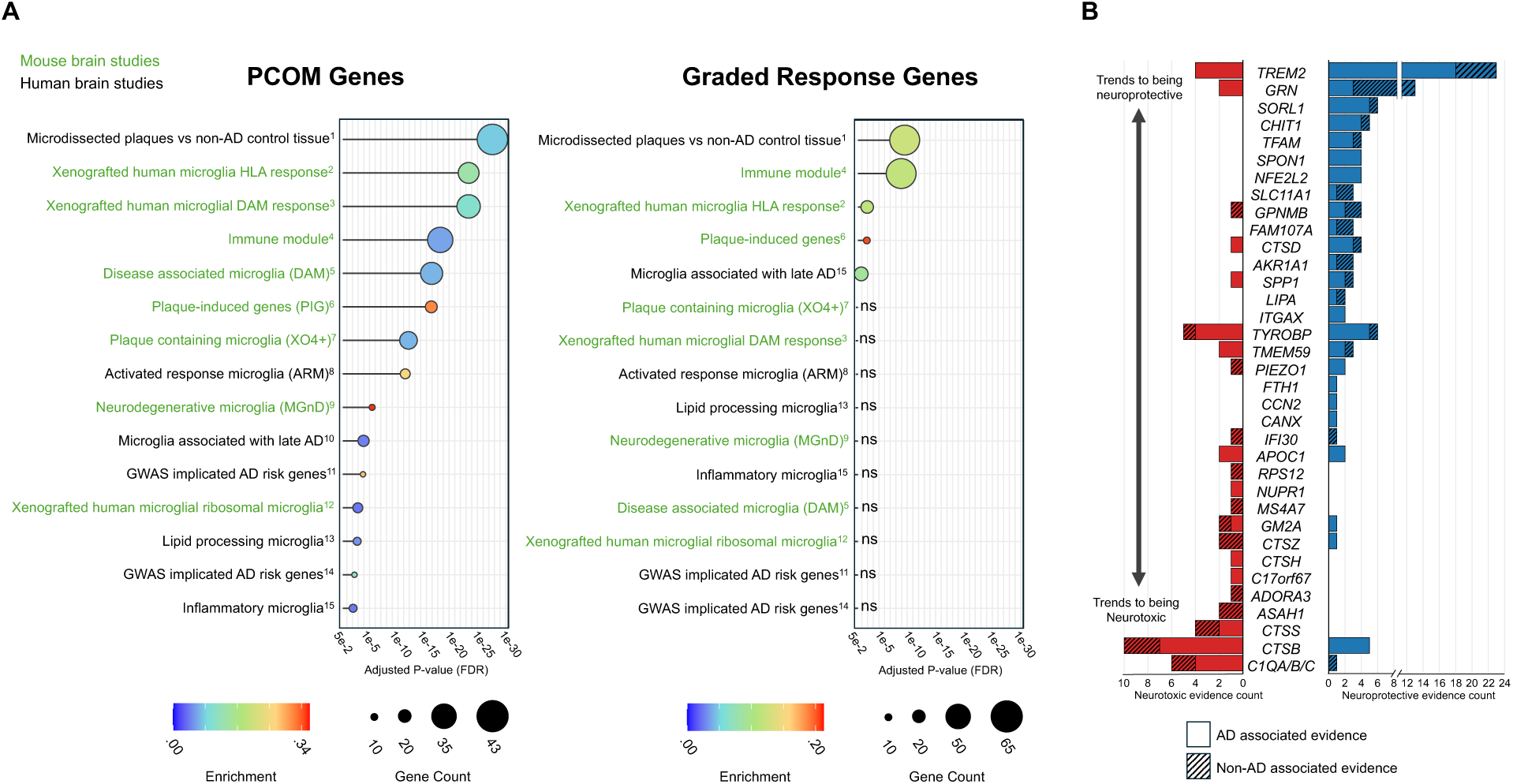
PCOM genes are significantly enriched for the DAM expression profile. (A) Literature enrichment analysis of PCOM and graded response gene categories.1: Das et al. 2024, 2: Mancuso et al. 2022, 3: Mancuso et al. 2022, 4: Salih et al. 2019, 5: Keren-Shaul et al. 2017, 6: Chen et al. 2020, 7: Grubman et al. 2021, 8: Frigerio et al. 2020, 9: Krasemann et al. 2017, 10: Sun et al. 2023, 11: Wightman et al. 2021, 12: Mancuso et al. 2022, 13: Sun et al. 2023 (MG4 cluster), 14: Bellenguez et al. 2022, 15: Sun et al. 2023 (MG2, MG8, & MG10 cluster). (B) Plot of evidence accumulation for PCOM genes with mechanistic studies supporting neuroprotective or neurotoxic roles in Alzheimer’s disease or other disease contexts. Statistical analysis: (A) P values were calculated using Fisher’s exact test and adjusted for multiple comparisons using Benjamini-Hochberg false discovery rate (FDR) method. Human cases (n = 26).

The upregulation of these gene modules is often broadly interpreted with a polarised view of either promoting neurotoxicity or conferring neuroprotection. However, emerging evidence, particularly from mechanistic studies of individual genes, reveals a potentially complicated picture. This is illustrated in the plot of evidence accumulation (PEA; Figure 3B), which summarises published experimental evidence supporting either neuroprotective or neurotoxic roles for key PCOM genes. While some genes, for example those encoding TREM2 and SPON1, show strong support for neuroprotective functions, others, such as CTSS, CTSB, and C1q investigations are more frequently associated with detrimental outcomes. These findings suggest that, while the majority of PCOM genes studied mechanistically appear to exert neuroprotective effects in AD, some may contribute to disease progression. Each gene and associated evidence can be found in Supplemental Table 4.

### Distinct gene modules are associated with disease stage and *APOE* status

To identify coordinated transcriptional responses, we performed weighted gene co-expression network analysis (WGCNA) on the spatial transcriptomic data (Figure 4A). Hierarchical clustering of gene expression profiles revealed five distinct gene modules. Correlation analysis between module eigengenes and covariates identified several modules with significant associations. The brown module, comprising 323 genes, exhibited higher eigengene values in plaque-contacting microglia and in APOE4 carriers, indicating that the genes within this module are, on average, more highly expressed under these conditions (Figure 4A). This module was centred around ribosomal proteins (*RPL* genes), which were highly interconnected with numerous immune-related genes, including *C3*, *C1QC*, and *B2M* (Figure 4Bi). Gene Ontology analysis confirmed the module’s involvement in both translational and immune-related processes (Figure 4Bii). To identify genes contributing most strongly to the brown module, module membership was stratified by both APOE status and plaque-contacting versus away microglia. Genes falling within the top 25% of module membership for both covariates were highlighted. This subset included several well-established AD-associated genes, such as the complement components *C1QC* and *C1QB*, the TREM2 signalling adaptor encoding *TYROBP*, and the transcription factor *SPI1* (Figure 4Biii), suggesting that these genes are closely associated with *APOE* status and plaque-contacting microglia.

**Figure 4.**
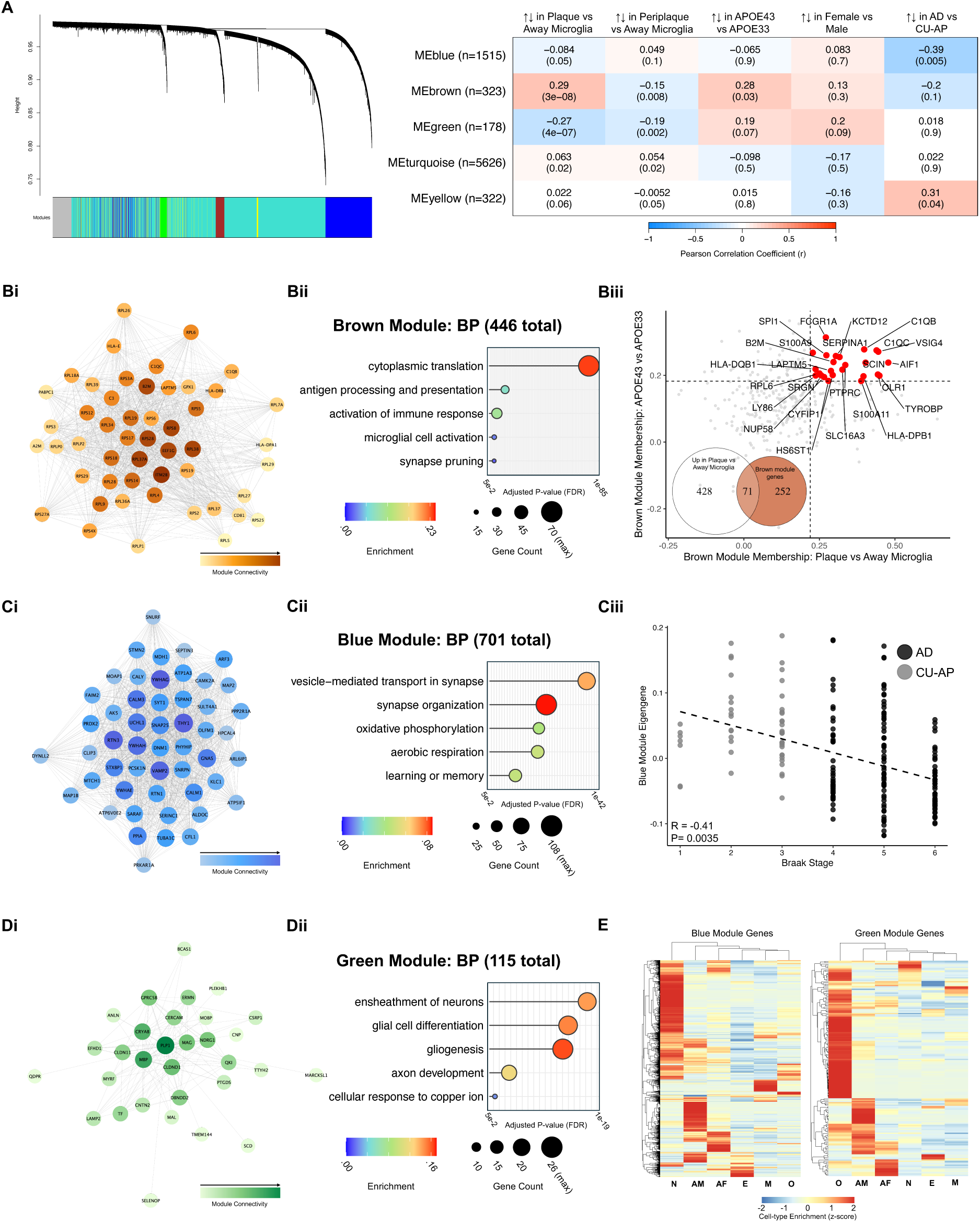
Distinct gene modules are associated with disease stage and APOE status. (A) Hierarchical clustering dendrogram (left) identifying five distinct gene modules, with corresponding module-trait correlations (right). (B) Brown module visualisation. (Bi) Network plot of the top genes ranked by connectivity. (Bii) Gene ontology analysis highlighting select enriched terms from the Biological Process database. (Biii) Module membership scores revealing genes in the top quartile (Q3) for both APOE status and plaque vs. away correlations, along with a Venn diagram showing overlap between brown module genes and differentially expressed genes (DEGs) in plaque vs. away microglia. (C) Blue module visualisation. (Ci) Network plot of the top genes ranked by connectivity. (Cii) Gene ontology analysis highlighting select enriched terms from the Biological Process database. (Ciii) Blue module expression across Braak stages. (D) Green module visualisation. (Di) Network plot of the top genes ranked by connectivity. (Dii) Gene ontology analysis highlighting select enriched terms from the Biological Process database. (E) Cell type enrichment analysis for the Blue and Green module genes based on cell-type expression values derived from Zhang et al. (2016). Statistical analysis: (A & Cii) Reported r values reflect Pearson correlations; statistical significance (p values) was assessed using linear mixed models to account for repeated measures. (Bi, Ci, & Di) Network plots show the top 50 genes per module based on connectivity. Modules with fewer than 50 genes exceeding the edge weight threshold (≥ 0.08) are shown in full. (Bii, Cii, & Dii) P values are adjusted for multiple comparisons using Benjamini-Hochberg false discovery rate (FDR) method. Human cases (n = 26).

The importance of the transcription factor encoding SPI1 within the brown module suggests it may play a key regulatory role in orchestrating the microglial transcriptional response to Aβ plaque pathology. To investigate this further, we first examined SPI1 expression directly. SPI1 was the only transcription factor gene showing both significantly increased expression and activity in the Plaque vs Away comparison (Figure S4A). Stratifying by APOE status revealed a significant interaction, with SPI1 expression being significantly higher in plaque-contacting microglia from APOE4 carriers compared to APOE3 (Figure S4B). To explore broader transcription factor activity, we then performed VIPER analysis which identified numerous transcription factors with significantly altered activity across the three AOI comparisons (Figure S4C). In this context, “activity” refers to the inferred regulatory influence of a transcription factor on its downstream targets, rather than its expression level, based on the DoRothEA regulon that catalogues experimentally supported transcription factor–target interactions. Among these, SPI1 exhibited the most significant increase in activity in the Plaque vs Away microglia comparison. Consistent with its contribution to APOE-associated brown module membership, overall SPI1 activity was also significantly elevated in APOE4 carriers (Figure S4Dii).

The blue module, comprising 1,515 genes, showed significantly reduced expression in AD compared to CU-AP individuals (Figure 4A). This module was centred by a highly interconnected network of synaptic genes, including *VAMP2*, *SNAP25*, and *SYT1*; mitochondrial-associated genes such as *MTCH1*, *ATP5IF1*, and *ATP1A3*; as well as genes encoding markers of dystrophic neurites, notably *RTN3* and *RTN1* (Figure 4Ci). Gene ontology analysis confirmed significant enrichment of synaptic and mitochondrial genes within the blue module (Figure 4Cii). Given that CU-AP individuals may represent a preclinical stage of AD, the observed reduction in blue module gene expression in the AD samples could reflect disease progression.

Supporting this, the blue module expression pattern showed a significant negative correlation with increasing Braak stage (Figure 4Ciii). On the other hand, the CU-AP group, despite being at low Braak stages (1-3), were predominantly APOE4/4 (5/7 individuals) and spread across the age range. So, this lower expression or other differences could reflect protective factors.

Finally, the green module of 178 genes showed reduced expression in both plaque-contacting and periplaque microglia vs. away regions (Figure 4A). This module was centred primarily on myelin proteins including, *PLP1*, *MBP* and *MAG* (Figure 4Di). Again, the myelin association was confirmed by gene ontology analysis (Figure 4Dii). The remaining yellow and turquoise modules lacked overt association with covariates but can be visualised in Figure S5.

The absence of immune associations in the blue and green modules was underscored by the lack of microglial specificity among their constituent genes (Figure 4E). This exemplifies the key limitation of GeoMx whereby transcripts from cells above or below the targeted microglia are also captured. While this contamination reduces microglial specificity, it simultaneously offers valuable insights into the responses of neighbouring cell populations. All modules and associated genes can be found in Supplemental Table 5. The expression profiles of all quality-controlled genes across covariates can be explored interactively at https://edwardslab.shinyapps.io/Humaneac/.

### Conserved microglial response to plaques in *App^NL-F^* mice and human Alzheimer’s disease

Our previous investigation into microglial-enriched gene expression towards Aβ plaque pathology explored the spatial specificity of plaque-induced gene expression (PIGs; Chen et al., 2020) in 18-month-old *App^NL-F^* mice (Wood et al., 2022). To compare the microglial responses from this AD mouse model data and human AD tissue, the transcriptomic data from 18-month-old *App^NL-F^* mice were similarly analysed across the whole transcriptome, mimicking the analysis pipeline of the human tissue analysis. Three identical pairwise comparisons were similarly made: plaque-contacting vs away microglia (Figure 5Ai), plaque-contacting vs periplaque microglia (Figure 5Aii), and periplaque vs away microglia (Figure 5Aiii).

**Figure 5.**
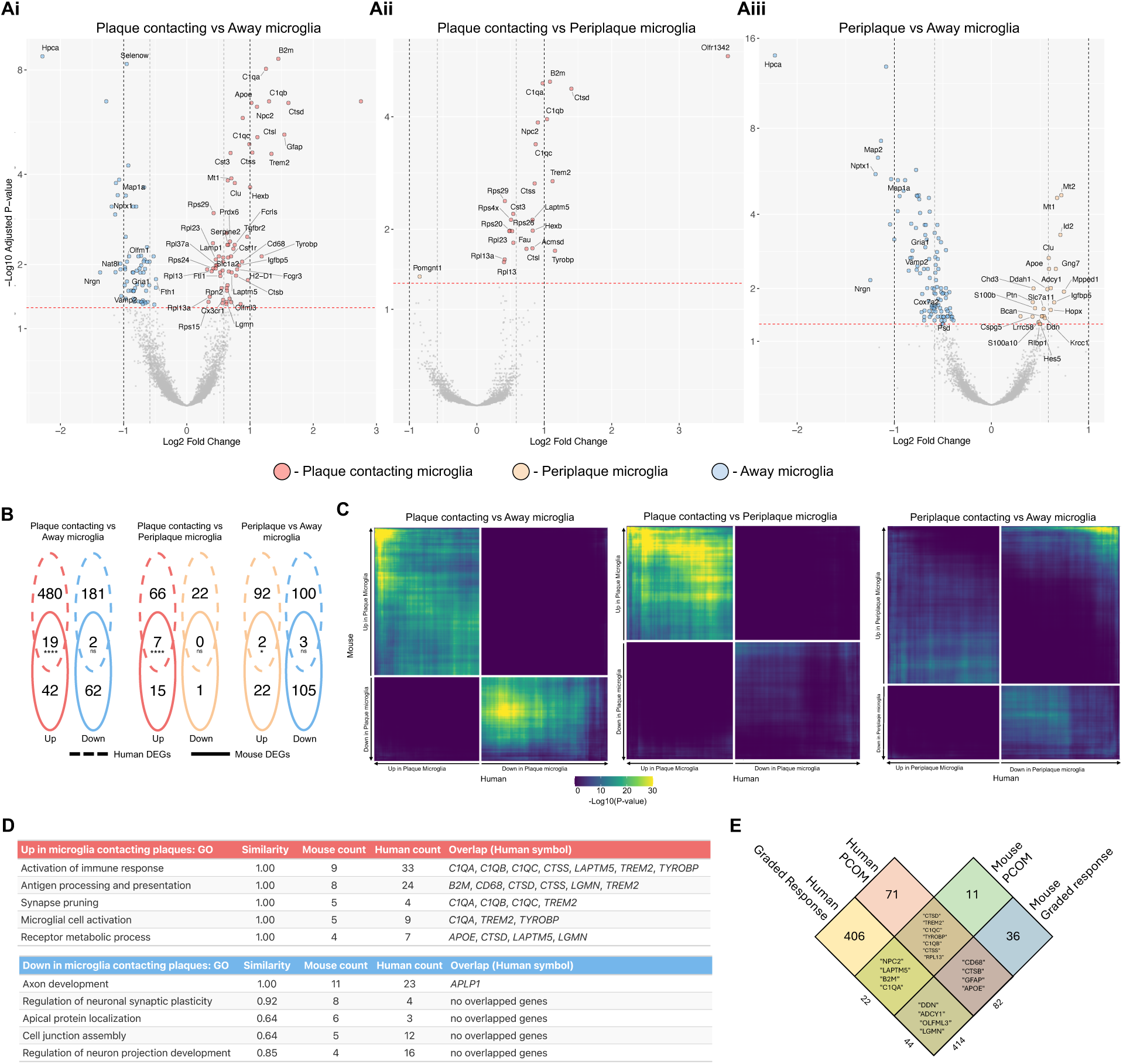
Conserved microglial response to plaques in *App^NL-F^* mice and human Alzheimer’s disease. (A) Volcano plots showing differential expression analyses from 18-month-old *App^NL-F^* mice for microglia in (Ai) Plaque vs Away, (Aii) Plaque vs Periplaque, and (Aiii) Periplaque vs Away regions. (B) Venn diagrams showing the overlap of significant differentially expressed genes (DEGs) across the three AOI comparisons in human tissue and *App^NL-F^* mouse analyses. (C) Rank–Rank Hypergeometric Overlap (RRHO) analysis comparing ranked gene expression changes in *App^NL-F^* and human microglia across matched AOI comparisons. Heatmaps display directional overlap of ranked gene expression changes. (D) Tables list GO:BP terms enriched among genes up-regulated (top) or down-regulated (bottom) in plaque-contacting microglia for similar GO terms significant in the human and mouse data, of which the human term is listed. Similarity is GO semantic similarity (0–1; 1.00 = identical term). Mouse count and Human count are the numbers of DEGs in each term. Overlap lists shared genes between the similar terms. (E) Overlap of human and mouse microglial gene expression for plaque-associated and graded response categories. Statistical analysis (A) (A) Statistical analysis using limma-voom with covariate adjustment and adjusted for repeated measures. P-values were adjusted for multiple comparisons using Benjamini-Hochberg false discovery rate (FDR) method. (B) P values were calculated using Fisher’s exact test. (C) Overlap was assessed using a hypergeometric test across all rank thresholds in a split-quadrant design. 18-month-old *App^NL-F^* mice (n=6) & human cases (n = 26).

While we aimed to replicate the experimental paradigm previously used in *App^NL-F^* mice closely, several adaptations were necessary to accommodate differences in the human AD tissue analysis. Specifically, due to staining efficacy, TMEM119 was used as the microglial marker in mouse tissue instead of IBA1 used in the human tissue analysis. An Aβ-specific antibody was employed to label plaques in mice, whereas an amyloid dye was used in human samples. Additionally, to account for interspecies differences in tissue scaling, periplaque microglia were defined as residing within 30 μm of a plaque in mice, compared to 50 μm in human tissue. Finally, it is important to note, that the earlier mouse study was hypothesis-driven and focused on a restricted subset of genes, meaning that its relatively small sample size was sufficient for targeted analyses but provides limited statistical power for the present whole-transcriptome comparison.

DEGs from each of the three AOI comparisons in the mouse analysis were compared with the corresponding orthologs in the human tissue dataset. This revealed significant overlap in differentially expressed genes upregulated in plaque-contacting versus away microglia, plaque-contacting versus periplaque microglia, and periplaque versus away microglia, illustrating a conserved spatial gene expression response between species (Figure 5B). Individual genes from these comparisons can be seen overlaid with the human tissue volcano plots in Figure S6.

Treating mouse DEGs as gene modules and mapping these to fold-change-ranked human genes using gene set enrichment analysis (GSEA) for each AOI comparison revealed a significant concordance in upregulated genes between mouse and human plaque-contacting microglia, particularly in the plaque vs. away and plaque vs. periplaque comparisons (Figure S7). In contrast, the periplaque vs. away comparison showed weaker enrichment, with downregulated DEGs from the mouse dataset unexpectedly aligning with upregulated genes in the human data, highlighting an incongruous interspecies result. Rank–rank hypergeometric overlap (RRHO) analysis was then performed to provide a threshold-free, directional assessment of overlap between human and mouse transcriptomic responses across each plaque-distance comparison (Figure 5C). In this method, genes are ranked by their expression change, and overlap is assessed iteratively at increasing thresholds (top 10, 20, 30, and so on) between the two species, with each quadrant of the plot representing concordance between genes that are either up- or down-regulated in each dataset. In both the plaque-contacting vs away and plaque-contacting vs periplaque comparisons, RRHO heatmaps revealed strong concordance between genes upregulated in mouse and human plaque-contacting microglia (top-left quadrant), indicating robust cross-species conservation of the plaque-induced transcriptional programme. Notably, when using this threshold-free approach, additional overlaps emerged among downregulated genes (bottom-right quadrant), though this concordance was more modest and did not extend to the most strongly downregulated transcripts. In contrast, the periplaque versus away comparison showed much weaker correspondence, consistent with earlier GSEA results, suggesting that while the plaque-contacting microglial response is well conserved between species, periplaque-responsive genes may not be. Next, we assessed gene ontology (Biological Process) enrichment for genes up- or down-regulated in plaque-contacting versus away microglia in both the human and mouse data. Terms were then matched across species by GO semantic similarity to highlight overlapping terms. As expected, immune-response processes were enriched in both species, with overlapping genes including the C1q subunits (*C1QA*, *C1QB*, *C1QC*), *TREM2*, and its signalling adaptor *TYROBP*. Despite limited overlap in the specific down-regulated genes, both human and mouse datasets converged on similar processes with synaptic and axonal-associated genes affected (Fig. 5D). The complete set of matched GO:BP terms enriched in plaque-contacting versus away microglia across human and mouse datasets is provided in Supplementary Table 6. Finally, to assess inter-species concordance in gene expression response patterns, genes were similarly categorised across species (Figure 2B). In both 18-month-old *App^NL-F^*mice and human AD tissue, key PCOM genes such as *TREM2* and its signalling partner *TYROBP*, complement components *C1QC* and *C1QB*, and the cathepsins *CTSS* and *CTSD*, were consistently categorised across species. The expression profiles of all quality-controlled genes for the 18-month-old *App^NL-F^* mouse data can be explored interactively at https://edwardslab.shinyapps.io/Humaneac/.

## Discussion

The presented study provides the first direct analysis of human gene expression specifically within microglia in contact with plaques. The results support the view that DAM and other reported AD-associated microglial states, whether identified in mouse models or human tissue, predominantly represent plaque-contacting populations. Furthermore, our coupled interspecies comparison, applying the same experimental paradigm in human AD tissue and mouse models, reveals a highly conserved upregulation response in plaque-contacting microglia across species. By contrast, genes changing between away and periplaque microglia show less interspecies congruence, indicating that plaque-contact–driven transcriptional changes are the most conserved component of the microglial response. Consistent with this, GO:BP analysis showed robust enrichment of immune-activation pathways. By contrast, although few down-regulated DEGs were shared across species, both datasets converged on synaptic and axonal terms, suggesting local synapse and axon loss in plaque-adjacent tissue, consistent with prior reports of plaque-proximal synaptic loss (Koffie et al., 2009; Masliah et al., 1989; Spires et al., 2005).

It is important to note that the human study employed a dye specific to fibrillar Aβ plaques, as Aβ antibodies were ineffective on human tissue following proteinase K digestion, whereas the mouse experiment used an Aβ antibody. As a result, a halo of non-fibrillar Aβ likely surrounds the fibrillar core identified by the dye used in the human study (Koffie et al., 2009; Wood et al., 2024). Consequently, the periplaque regions in the human samples may contain higher concentrations of Aβ than the periplaque regions in the mouse experiment, where both fibrillar and non-fibrillar forms were detected. This methodological difference likely contributed to the observed discrepancies in plaque or peri-plaque-associated gene expression between the two species. Another important difference to consider is that the mice are more equivalent to the CU group in the human comparison as they have an increasing plaque load but have not developed neurofibrillary tangles. Consequently, it is perhaps not surprising that it is particularly the directly plaque-associated changes that show the strongest similarity with the human data.

The distribution of ligands driving these spatial gene-expression responses is likely to differ relative to plaques. Plaque-contact specific responses (PCOM genes) may be triggered by ligands enriched within plaques themselves, such as lipid species, damage-associated molecules like phosphatidylserine, or plaque-associated proteins such as APOE, all of which are recognised TREM2 ligands (Atagi et al., 2015; Hardy and Escott-Price, 2019; Scott-Hewitt et al., 2020; Wang et al., 2015b). TREM2’s role in regulating the microglial transcriptional response to amyloid plaques has been consistently demonstrated (Grubman et al., 2021; Keren-Shaul et al., 2017). In our previous mouse study, *App^NL-F^* mice carrying the *Trem2^R47H^*mutation confirmed that functional TREM2 is critical for controlling the transcriptional response of microglia in direct contact with plaques (Wood et al., 2022). While the present study cannot directly determine whether these PCOM genes in human tissue are TREM2-dependent, the selective upregulation of *TREM2* and signalling partner *TYROBP* in plaque-contacting microglia, together with the enrichment of known TREM2-dependent genes such as *C1QB*, *C1QC*, and *CTSS*, is consistent with the possibility that TREM2 also plays an important role in controlling this response in humans.

Within the PCOM category, the genes enriched ontology terms related to phagocytosis, lysosomal function, and lipid metabolism. As well as *TREM2,* examples include: complement components *C1QB* and *C1QC*; lipid transporters such as *APOE* and *APOC1;* lysosomal regulators including *GRN*, *CD68 and LIPA* as well as multiple cathepsins (*CTSD*, *CTSB*, *CTSH*, *CTSZ*). These findings suggest that plaque-contacting microglia are actively engaged in ingesting, degrading, and storing plaque-associated material, particularly lipids. Such phagocytic activity at plaques has been extensively documented, with synapses and Aβ identified as key targets (Condello et al., 2015; D’Andrea et al., 2004; Tzioras et al., 2023). In parallel, consistent with their interaction with lipids, lipid droplet accumulation in microglia surrounding plaques has been observed (Claes et al., 2021; Prakash et al., 2024).

Genes that increased gradually toward plaques enriched terms associated with cell communication and migration, such as *CCL13*, *MYD88*, and *CD74*, as well as pro-inflammatory indicators such as *IL13*, the TNF receptor *TNFRSF1B* and the inflammasome-associated genes *P2RX7* and *PYCARD*. This pattern is consistent with an activation by diffusible factors that form a concentration gradient from the plaque, providing danger signals that potentially prime the microglial response (Wang et al., 2015a).

Network analysis revealed distinct co-expression modules with strong associations to both plaque proximity and other covariates. The brown module showed substantial overlap with immune DEGs upregulated in plaque-contacting microglia, but the network was centred on genes encoding translation-associated machinery.

Increases in ribosomal proteins have also been reported alongside immune gene upregulation in microglia during ageing (Ximerakis et al., 2019). Consistent with this, ‘ribosome’ was among the most significantly enriched KEGG pathways in a transcriptomic study of microglia containing ingested plaque aggregates, driven by increased expression of *RPL* genes, several of which overlapped with hub genes in the brown module (Grubman et al., 2021). This ribosomal protein hub is therefore likely to support targeted local protein synthesis required for the expression of connected immune-associated genes. This module also showed significantly higher activity and expression in APOE4 carriers compared to APOE3, potentially contributing to the elevated risk associated with APOE4. Alternatively, if this module reflects a protective response, the increased activity observed in APOE4 carriers may instead represent a compensatory reaction to prior APOE4-related toxicity, rather than a direct pathogenic effect. Notably, SPI1 exhibited high module membership for both APOE status and microglial responses to plaques implying that its increased expression in APOE4 carriers could relate to the increased risk of AD. Overexpression of SPI1 has previously been linked to reduced microglial viability and heightened expression of pro-inflammatory cytokines (Jones et al., 2021; Pimenova et al., 2021; Rustenhoven et al., 2018). Similarly, genetic evidence indicates that reduced expression of the *SPI1*-encoded transcription factor PU.1 is associated with a lower risk of AD (Huang et al., 2017). While PU.1 has been shown to regulate *APOE* expression (Pimenova et al., 2021) and the APOE4 genotype has been reported to promote a pro-inflammatory microglial phenotype in AD (Dias et al., 2025), there is limited evidence directly connecting this effect to *SPI1*.

The blue module comprised mitochondrial and synaptic genes whose expression declined with increasing Braak stage. A proteomic meta-analysis by Askenazi et al. (2023) reported a similar module of mitochondrial and synaptic proteins that decreased across stages of cognitive decline. Notably, the overlap between these datasets highlights enrichment for electron transport chain components, with complex I subunits showing particular vulnerability. Moreover, PET scans of complex I binding availability have shown a close relationship with Tau load over plaque load, suggesting a potential interaction between tau pathology and electron transport chain function (Terada et al., 2021). Consistent with this, wild-type Tau has been shown to interact with multiple mitochondrial electron transport chain components.

Interestingly synaptic vesicle fusion components were also implicated. These physiological interactions are diminished by frontotemporal dementia–causing Tau mutations, leading to impaired mitochondrial bioenergetics (Tracy et al., 2022). This is consistent with the changes suggested by the Blue module over Braak stages, suggesting these human-specific changes are Tau-related.

### Conclusion

We have demonstrated that, in human tissue, like in mice, upregulation of many of the genes that have been reported to respond to plaque load is dependent on direct contact with plaques. This includes *TREM2* and pathways controlled by TREM2, possibly under the SPI1 promotor. The concurrence of genes upregulated in microglia touching plaques in the two species supports the relevance of the mouse model in relation to the microglial response to plaques. In contrast, the differences between the species seen in areas further from plaques likely relate to the stage of disease progression, with additional changes occurring as neurofibrillary tangles and neurodegeneration proceed, pathologies that are not occurring in the mouse. We present a resource in which expression of individual genes of interest can be accessed in relation to plaques and compared across species.

## Contributions

Conceptualisation: J.I.W., F.A.E.

Methodology: J.I.W., G.P, S.D., D.G., M.B., D.S.

Experiments: J.I.W.,

Analysis: J.I.W., P.R., A.B., S.C.

Visualisations: J.I.W., P.R. Original Draft: J.I.W., F.A.E.

Review and editing: J.I.W., G.P, P.R., S.D., S.C., A.B., D.G., M.B., D.S, J.Hardy,

J.Hanrieder. F.A.E.

Supervision: D.M.C., D.S., J. Hanrieder, F.A.E, Funding: J. Hardy, D.M.C., J. Hanrieder, F.A.E.

## Acknowledgements

This project was funded by The Cure Alzheimer’s fund (F.A.E. and J.H.). J.I.W. is funded by the Swedish Alzheimer’s foundation. The initial work carried out by G.P. was supported through a Genetics Society-funded internship. J.H. is supported by the Dolby Foundation and by the NIHR University College London Hospitals Biomedical Research Centre. J.H. and D.S. are supported by the UK Dementia Research Institute through UK DRI Ltd, principally funded by the Medical Research Council (MRC). P.R. was funded by Ecole Normale Supérieure Paris-Saclay.

