## Supplemental Figures for "Spatially Resolved Microglial Expression Around Aβ Plaques in Human Alzheimer’s Disease Tissue"

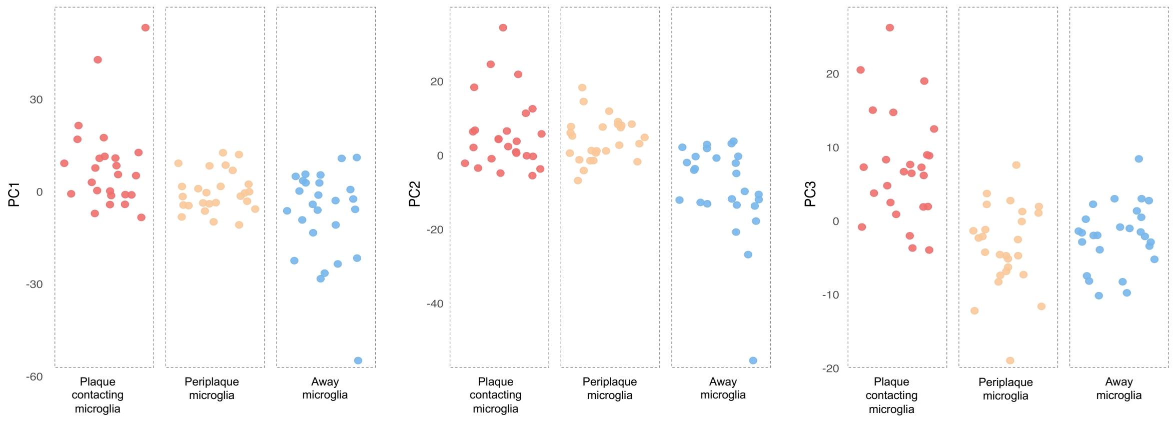


**Figure S1. Principal component analysis of gene expression across AOIs**
PCA was performed on voom-transformed $\log_2$CPM values, corrected for batch, case, and covariates (APOE genotype, sex, and clinical diagnosis).

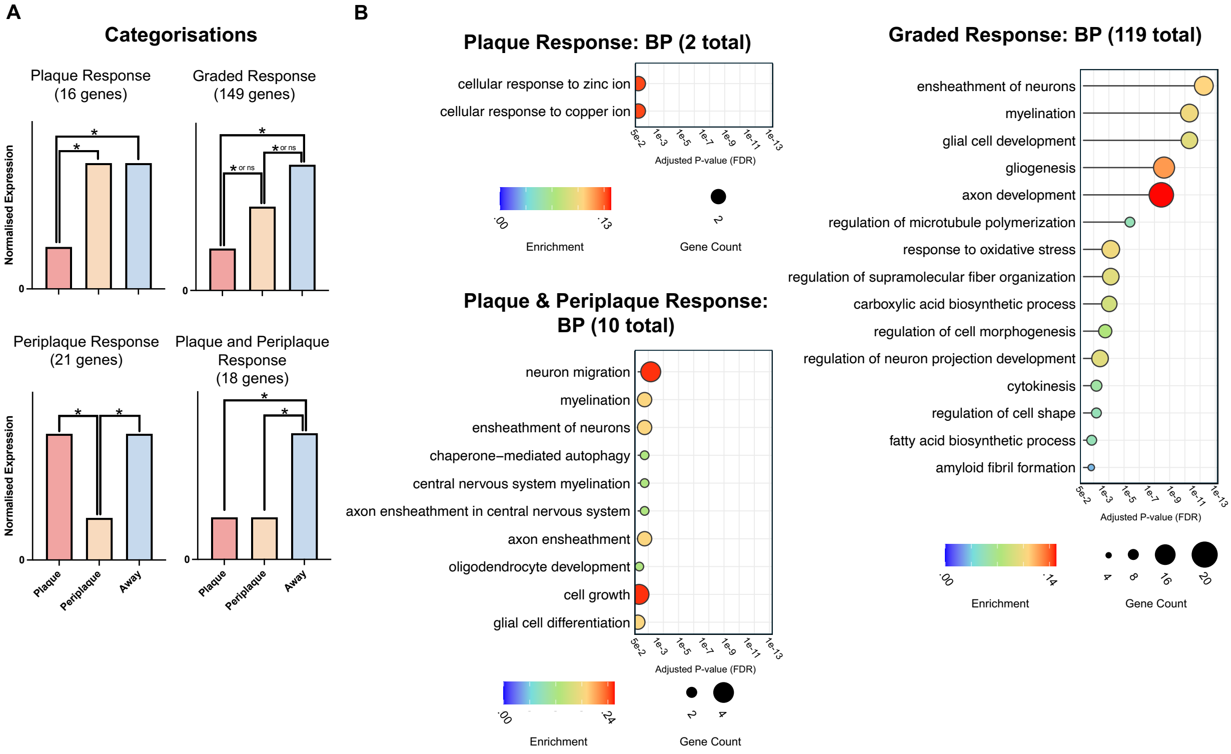


**Figure S2. Down regulated gene categories with associated gene ontology analysis.** (A) Representative categories of downregulated genes illustrating different gene expression response to plaques, with example graphs for each category. (B) GO Biological Process enrichment analysis of genes downregulated in microglia across three response types: specifically upon plaque contact (plaque response), showing a gradual decrease with proximity to the plaque (graded response), and downregulated in both plaque and periplaque regions (plaque and periplaque response). Selected enriched terms are shown, with the total number of significant terms identified in each group indicated in brackets. (C) Gene ontology enrichment was performed using the enrichGO function from the clusterProfiler package. P values were calculated using a hypergeometric test and adjusted for multiple comparisons using the Benjamini-Hochberg false discovery rate (FDR) method. Human cases (n = 26).

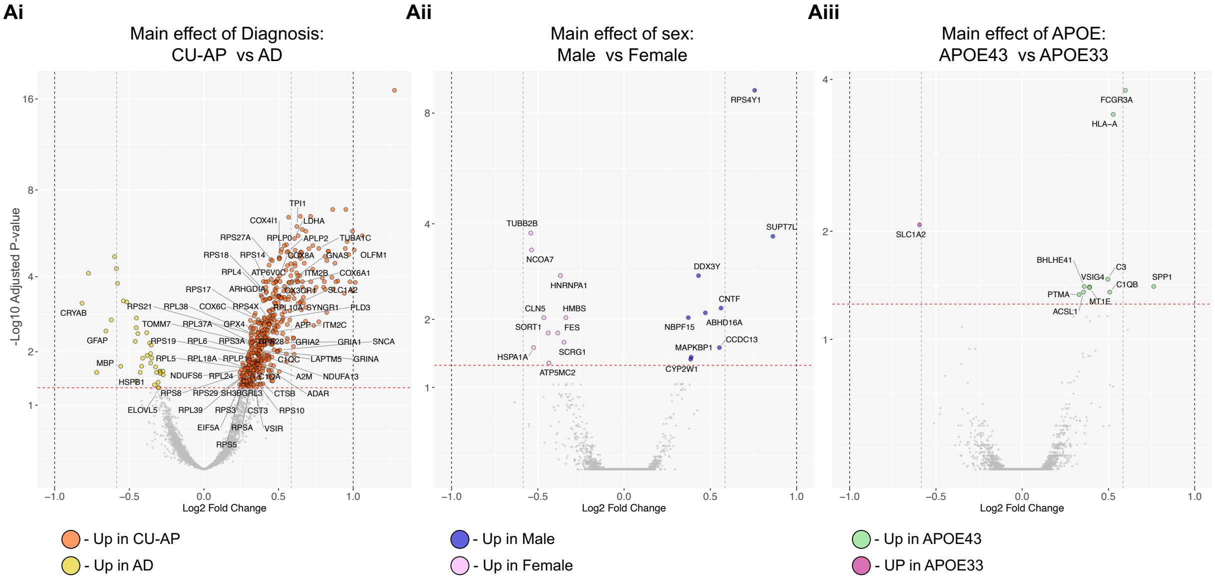


**Figure S3. Main effects of covariates on gene expression across all ROIs.**
(A) Volcano plots showing genes differentially expressed genes with a (Ai) main effect of diagnosis (CU-AP individuals and individuals with AD), (Aii) main effect of sex (male vs female), and (Aiii) main effect of APOE genotype (APOE44 vs APOE33). Differential expression was performed using linear modelling with empirical Bayes moderation (limma-voom) accounting for repeated measures by subject. P-values were adjusted for multiple comparisons using Benjamini-Hochberg false discovery rate (FDR) method. Human cases (n = 26).

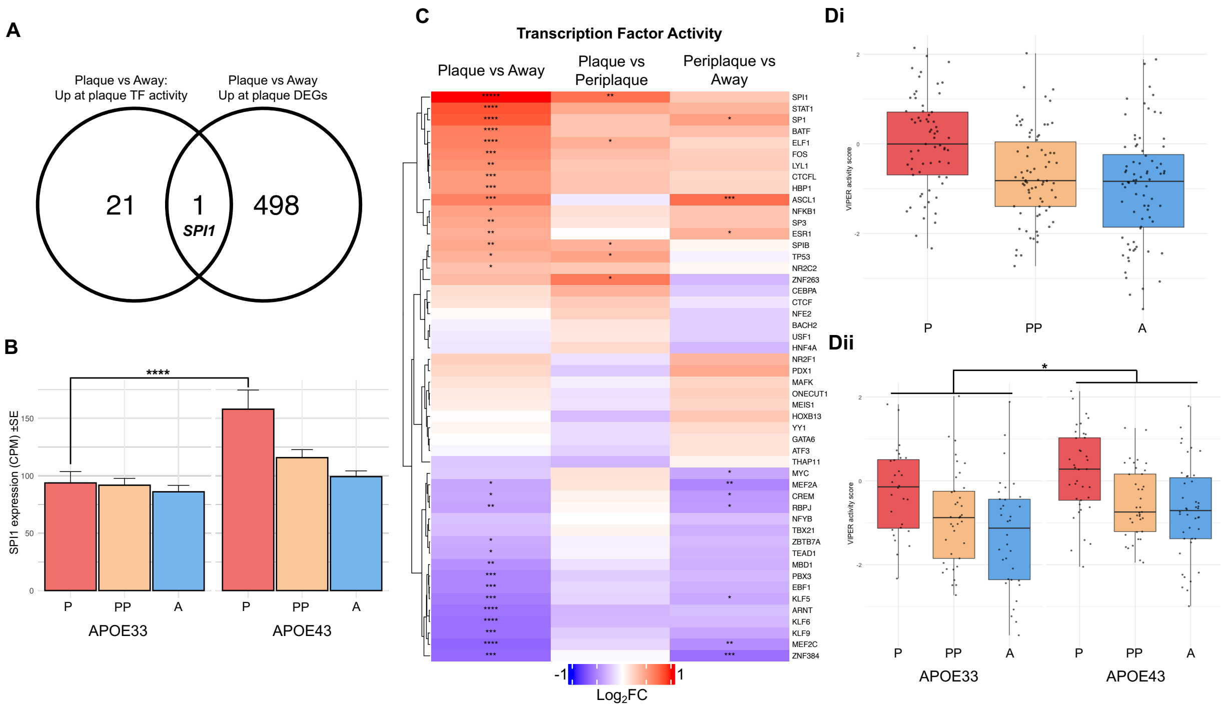


**Figure S4. Transcription factor activity is altered in plaque-contacting microglia, with SPI1 activity and expression modulated by APOE genotype.**

(A) Venn diagram showing the overlap between transcription factors with significantly increased activity (VIPER-inferred) and upregulated differentially expressed genes (DEGs) in plaque vs. away regions. (B) *SPI1* gene expression (CPM, batch-adjusted) across AOIs and stratified by APOE status activity (LMM: main effects: AOI type P<0.0001; *APOE* status P<0.01; interaction P<0.01).(C) Heatmap showing transcription factor activity scores calculated by the VIPER R package and based on the DoRothEA transcription factor database. Comparisons were made between plaque-contacting (P), periplaque (PP), and away from plaque. (Di) Viper-calculated SPI1 activity across P, PP and AOIs across all samples. (Dii) Stratification by *APOE* status of SPI1 activity (LMM: main effects: AOI type P<0.001; *APOE* status P<0.05; interaction P>0.1). Statistical analysis: (B & Dii) linear mixed model with AOI type & APOE status, and accounting for subject repeated measures. (B) Followed by Tukey-adjusted pairwise comparisons. (C) P-values for differential transcription factor activity were derived from linear modelling using limma, applied to VIPER-calculated activity scores and FDR-adjusted using the Benjamini-Hochberg method. (B, Bi, & Bii) data plotted as mean + SEM. Significance levels: ****P<0.0001; ***P<0.001; **P<0.01; *P<0.05. Human cases (n = 26 from n=15 APOE34, n=11 APOE33).

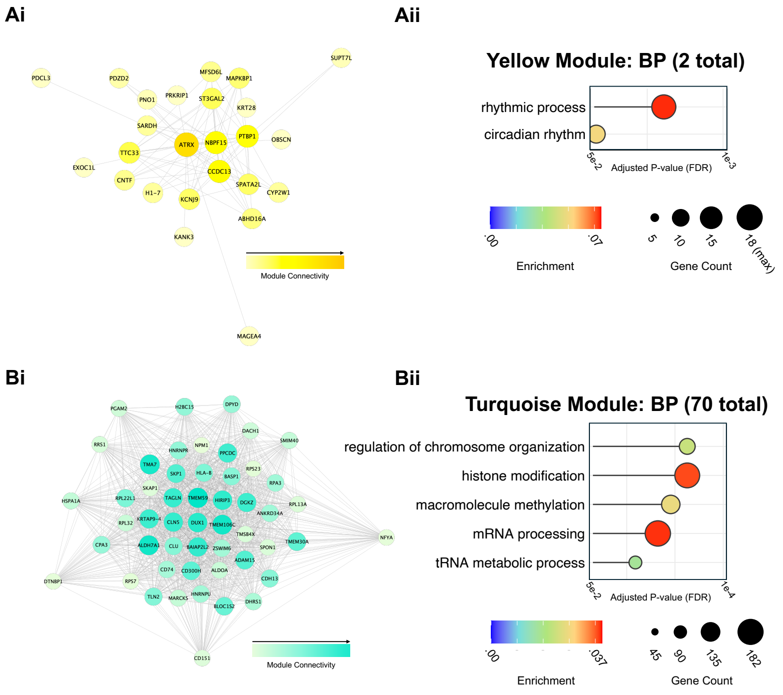


**Figure S5. Visualisation of the remaining Yellow and Turquoise WCGNA modules**
(A) Yellow module visualisation. (Ai) Network plot showing the top genes ranked by connectivity. (Aii) Gene ontology analysis highlighting enriched terms from the Biological Process database. (B) Turquoise module visualisation. (Bi) Network plot of the top genes based on connectivity. (Bii) Gene ontology analysis highlighting select enriched terms from the Biological Process database. Statistical analysis: (Ai & Bi) Network plots display all genes above the edge weight threshold (≥ 0.08). (Aii & Bii) Gene ontology terms were selected from the Biological Process category; p values are FDR-adjusted using the Benjamini-Hochberg method. Human cases (n = 26).

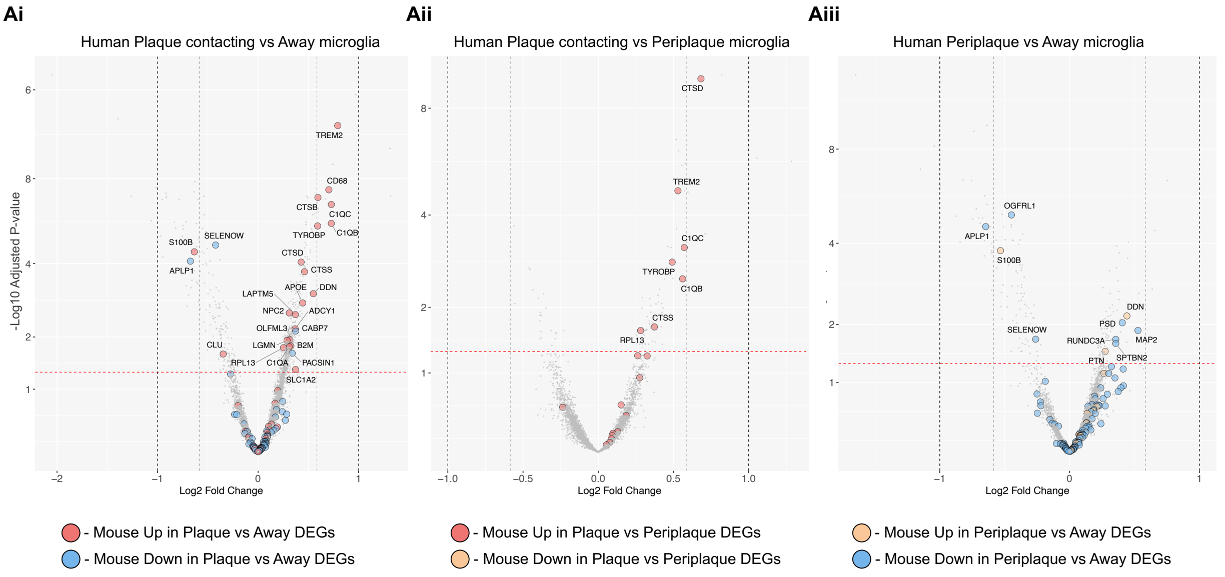


**Figure S6. Mouse DEGs for relative comparisons labelled on human AOI volcano plots.** (A) Volcano plots of human microglial differential expression analyses for (Ai) Plaque-contacting vs Away, (Aii) Plaque-contacting vs Periplaque, and(Aiii) Periplaque vs Away AOIs. Genes previously identified as significantly upregulated or downregulated in corresponding AOI comparisons in *App$^{NL\text{-}F}$* mice are overlaid and labelled on the human datasets following conversion to human orthologues. P-values were adjusted for multiple comparisons using Benjamini-Hochberg false discovery rate (FDR) method. 18-month-old *App$^{NL\text{-}F}$* mice (n=6) & human cases (n = 26).

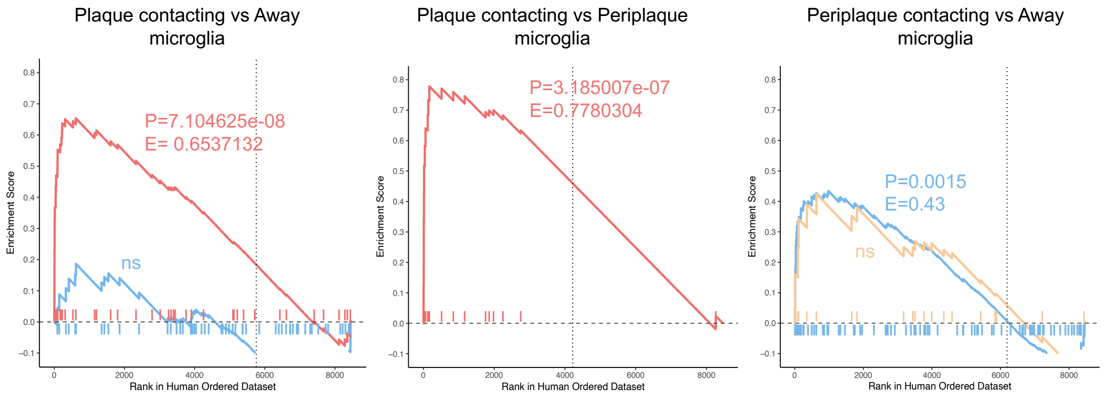


**Figure S7. GSEA of *App*$^{NL\text{-}F}$ mouse DEGs against fold-change–ranked human AOI lists.** Gene set enrichment analysis (GSEA) of *App*$^{NL\text{-}F}$ mouse differentially DEGs from the three AOI comparisons projected onto fold change–ranked human gene lists from the corresponding AOI comparisons. Enrichment scores were calculated using the GSEA running-sum statistic, with statistical significance assessed by permutation testing. 18-month-old *App*$^{NL\text{-}F}$ mice (n=6) & human cases (n = 26).
